# Prediction of specific TCR-peptide binding from large dictionaries of TCR-peptide pairs

**DOI:** 10.1101/650861

**Authors:** Ido Springer, Hanan Besser, Nili Tickotsky-Moskovitz, Shirit Dvorkin, Yoram Louzoun

## Abstract

Current sequencing methods allow for detailed samples of T cell receptors (TCR) repertoires. To determine from a repertoire whether its host had been exposed to a target, computational tools that predict TCR-epitope binding are required. Currents tools are based on conserved motifs and are applied to peptides with many known binding TCRs.

Given any TCR and peptide, we employ new NLP-based methods to predict whether they bind. We combined large-scale TCR-peptide dictionaries with deep learning methods to produce ERGO (pEptide tcR matchinG predictiOn), a highly specific and generic TCR-peptide binding predictor.

A set of standard tests are defined for the performance of peptide-TCR binding, including the detection of TCRs binding to a given peptide/antigen, choosing among a set of candidate peptides for a given TCR and determining whether any pair of TCR-peptide bind. ERGO significantly outperforms current methods in these tests even when not trained specifically for each test.

The software implementation and data sets are available at https://github.com/louzounlab/ERGO

## Introduction

T lymphocytes (T cells) are pivotal in the cellular immune response^1,2^. The immense diversity of the T-cell receptor (TCR) enables specific antigen recognition^3,4^. Successful recognition of antigenic peptides bound to major histocompatibility complexes (pMHCs) requires specific binding of the TCR to these complexes^5–7^, which in turn modulates the cell’s fitness, clonal expansion, and acquisition of effector properties^7^. The affinity of a TCR for a given peptide epitope and the specificity of the binding are governed by the heterodimeric αβ T-cell receptors^2^. While both chains have been reported to be important to affect binding, we show here that the TCR’s binding to target MHC-peptide can be determined for many TCR-peptide pairs with high accuracy using the β-chain only.

Within the TCRβ chain, the complementarity-determining region 1 (CDR1) and CDR2 loops of the TCR contact the MHC alpha-helices while the hypervariable CDR3 regions interact mainly with the peptide^1,2^. In both TCRα and TCRβ chains, CDR3 loops have the highest sequence diversity and are the principal determinants of receptor binding specificity.

Following specific binding of T cell receptors to viral and bacterial-derived peptides bound to MHC^5^, or from neo-antigens^8–10^, the appropriate T cells expand, resulting in the increased frequency of T cells carrying such receptors. Recently, high-throughput DNA sequencing has enabled large-scale characterization of TCR sequences, producing detailed T cell repertoire (Rep-Seq)^11^. Expanded clones are more likely to be repeatedly sampled in Rep-Seq than non-expanded clones and can serve as biomarkers for previous or current exposures to their cognate target, especially if the sample is enriched for mature or activated cells, or if strict filtering on sampled clone size is applied. Tools for the precise distinction between TCRs binding distinct targets are required to use T cell repertoires as systemic biomarkers (often referred to as “reading the repertoire”).

A direct approach for using TCR Rep-Seq as biomarkers has been proposed by Emerson et al.^12^ and similar approaches^13^ who detected patients that have CMV based on their full repertoire and the choice of TCRs that differ between CMV positive and negative patients. This approach is based on the presence of highly specific and repetitively observed public TCR in the response of different hosts to the same peptide (often denoted public clones, although the definition of such clones varies among authors^14^). Such an approach requires extensive repertoire sequencing for every condition tested.

In contrast, many TCR responses are characterized by a high level of cross-reactivity with single TCRs binding a large number of MHC-bound peptides, and single peptides binding a large number of TCRs^15,16^. TCRs binding the same MHC-peptide may share similarities but possess different CDR3 sequences. Thus, while for public clones the task of deciphering the relation between a peptide and the TCR binding is based on tallying the candidate public TCR, for most highly cross-reactive TCRs, a probabilistic approach is required.

Important steps have been made in this direction by Glanville et al.^4^ and Dash et al.^17^, who detected the clear signature of short amino acid motifs in the CDR3 region of TCRβ and TCRα in response to specific peptides presented by specific MHC molecules. This work was then extended by recent efforts that combined these motifs with machine learning to predict peptide-specific TCRs vs. naïve TCRs, using Gaussian Processes^18^, or predicting TCR-epitope binding with Convolutional Neural Networks^19,20^. These methods significantly outperform random classification in the distinction of TCR binding a specific peptide and random TCRs.

The next required step for using the repertoire to develop specific biomarkers would be to distinguish between TCR binding different peptides. An essential step in the development of high precision predictors is the standardization of the comparison methods. We propose three different tests, each with different outcomes, as the standard method to estimate such predictions (Figure 1A):

**Figure 1:**
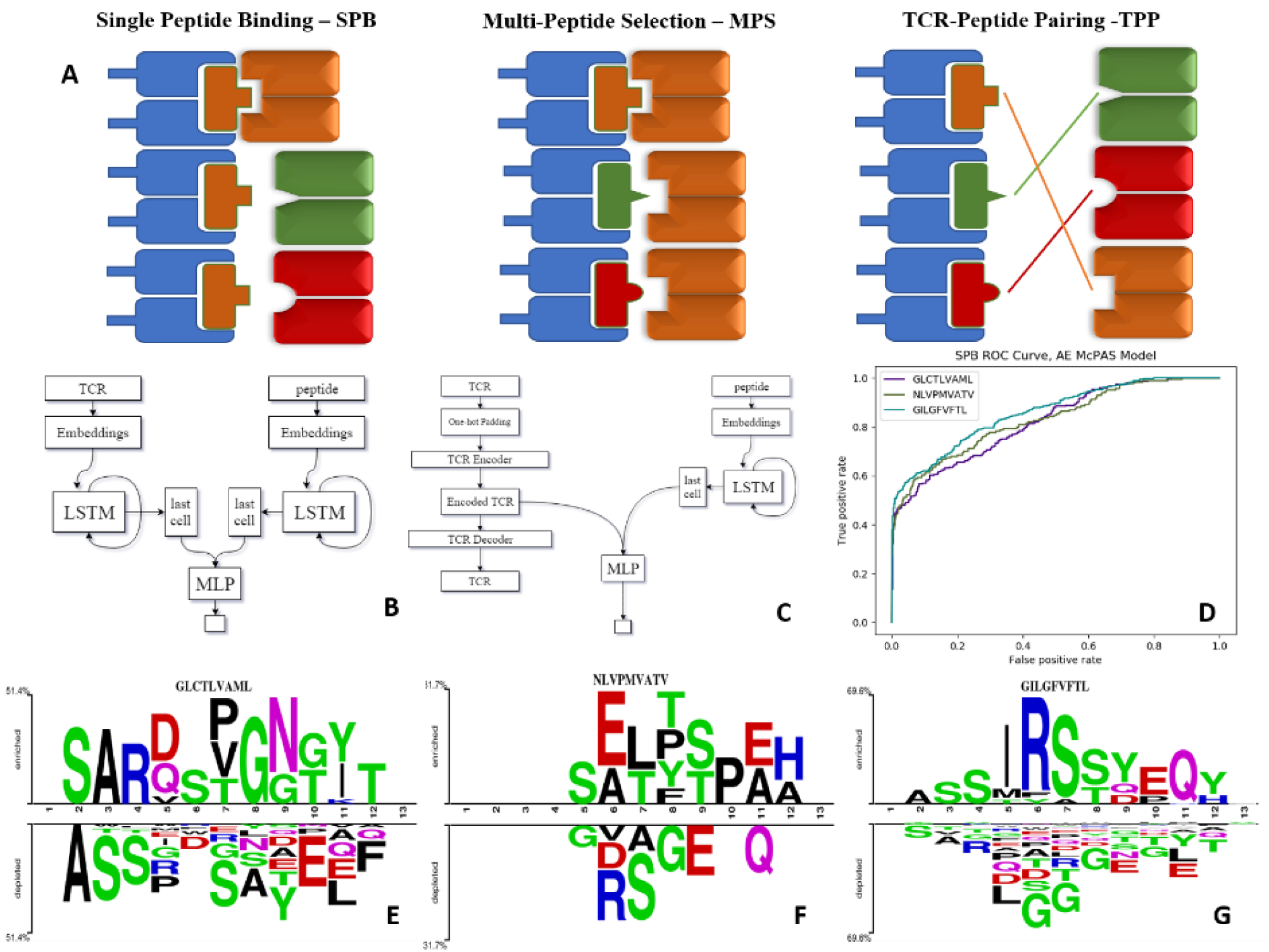
A) Illustration of the tests we suggest for evaluating the model performance as explained – Single Peptide Binding (SPB), Multi-Peptide Selection (MPS) and TCR-Peptide pairing (TPP). B) LSTM based model architecture. C) Autoencoder based model architecture. D) ROC curve of autoencoder based model SPB performance on 3 human peptides from Dash et al. dataset. E-G) Comparison of amino acids of CDR3 beta sequences of TCRs binding Dash et al. peptides vs. TCRs which do not bind these peptides, in McPAS database. (logos were created with Two-Sample-Logos); The height of symbols within the stack in each logo indicates the relative frequency of each amino acid at that position. Only amino acids whose distribution differ significantly between the two sets are shown, and only 13 length TCRs were compared.

- Single Peptide Binding – SPB. Testing whether an unknown TCR binds a predefined target, using (as training information) TCRs known to bind to this target^17,18^. The outcome of such a prediction would be the Area Under Curve (AUC) for the binding of an unseen TCR to this target.
- Multi-Peptide Selection – MPS. Given a set of predefined peptides, predict which of those will be bound by a new TCR. The outcome of that would be the accuracy of the choice as a function of the number of candidate peptides.
- TCR-Peptide Pairing -TPP. This is the most complex test. Given a large set of peptides and TCRs, test whether a randomly chosen TCR binds a randomly chosen peptide. This test can be further divided into three possibilities, based on the information used in the training stage: Previously observed peptide and TCR that are not known to pair (TPP-I), same with a previousely unobserved TCR (TPP-II) and finally the same with both a previousely unobserved peptide and a previousely unobserved TCR (TPP-III).

We propose these different tests as standard measures for the quality of TCR-peptide binding predictions. The TPP task is often addressed in Natural Language Processing (NLP) using recurrent neural networks (RNN)^21^. Long short-term memory (LSTM) networks are common type of RNN^22^. We employed LSTMs that produce an encoding of the varying TCR and peptide into constant length real-valued encodings and created ERGO (pEptide tcR matchinG predictiOn). We show that ERGO significantly outperforms existing methods in both single peptide binding prediction and multi-peptide selection, although it is only trained for the TPP task.

## Results

### ERGO outline

Target peptides and TCRs have different generation mechanisms (TCRs through VDJ recombination and junctional diversity^11^, and peptides through antigen generation, trafficking, processing and MHC binding^23^). Thus, ERGO uses different parallel encoders. At the broad level, we encode the CDR3 of each TCR and each peptide into numerical vectors. The encoded CDR3 and peptide are concatenated and used as an input to a feed-forward neural network (FFN), which should output 1 if the TCR and peptide bind and 0 otherwise (Figures 1B,1C). All models are trained to predict only the TPP-I task but are tested on all tasks. At this stage, the MHC was not included, since it did not contribute significantly to prediction accuracy in the current formalism.

For the peptides, we first use an initial random embedding and translated each amino acid (AA) into a 10-dimensional Embedding Vector. Changing the encoding dimension did not significantly change the obtained accuracy. In order to merge the encoding vectors of each position into a single vector representing the peptide, each vector was used as input to a LSTM. We used the last output of the LSTM as the encoding of the whole sequence. The embedding matrix values, the weights of the LSTM and the weights of the FFN were trained simultaneously. For the TCR encoding, we either used a similar approach or an autoencoder (AE) (See Supp. Mat. Methods and Figure 1C).

These models were trained on two large datasets of published TCR binding specific peptides^24,25^. McPAS-TCR^24^ is a manually curated database of TCR sequences associated with various pathologies and antigens based on published literature, with more than 20,000 TCRβ sequences matching over 300 unique epitope peptides. These TCRs are associated with various pathologic conditions (including pathogen infections, cancer, and autoimmunity) and their respective antigens in humans and in mice. VDJdb^25^ is an open, comprehensive database of over 40,000 TCR sequences and over 200 cognate epitopes and the restricting MHC allotype acquired by manual processing of published studies. For each TCR-peptide pair, a record confidence score was computed.

### ERGO can predict TCR binding to specific epitopes or antigens (SPB task)

ERGO was trained to solve the TPP-I problem (pairing TCR and peptide) on the two datasets above, and then tested on all three mentioned tests. To test the performance of ERGO on the SPB task (detecting whether a previousely unseen TCR binds a known peptide), we analyzed the five most frequent peptides in each datasets and tested the possibility of detecting whether a randomly selected TCR binds the peptide. The AUC for the binary classifications ranged between 0.695 and 0.980 (Table 1). The results are not sensitive to the number of TCRs reported for the peptides, and all peptides with more than 50 reported TCRs had similar values (Table S4).

**Table 1:** Comparison between the different versions of the ERGO classifier (AE (Autoencoder) vs. LSTM and McPAS vs VDJdb) for the SPB task. The five most frequent peptides in each database are shown. Other less frequent peptides SPB results are in the Supp. Mat. The values are the AUC over the test set of a previousely unseen TCR for this peptide.

| Peptide | McPAS |  | Peptide | VDJdb |  |
| --- | --- | --- | --- | --- | --- |
|  | AE | LSTM |  | AE | LSTM |
| LPRRSGAAGA | 0.772 | 0.760 | KLGGALQAK | 0.695 | 0.731 |
| GILGFVFTL | 0.843 | 0.832 | GILGFVFTL | 0.820 | 0.817 |
| NLVPMVATV | 0.835 | 0.821 | NLVPMVATV | 0.665 | 0.686 |
| GLCTLVAML | 0.803 | 0.816 | AVFDRKSDAK | 0.676 | 0.695 |
| SSYRRPVGI | 0.969 | 0.980 | RAKFKQLL | 0.828 | 0.825 |

To compare ERGO to current approaches, we tested its performance on current tools that predict TCR-peptide binding. We first compared it to the work of Jokinen et al.^18^ who compared TCRs found by Dash et al.^17^ to bind three human epitopes and seven mice epitopes with TCRs from VDJdb database^25^, which bind additional 22 epitopes. These peptide-TCR pairs were compared with naïve TCRs not expected to recognize the epitopes. Jokinen et al. evaluated the TCRGP model using leave-one-subject-out cross-validation (LOSO). The TCRGP model was trained with all subjects but one at a time and tested on the last. In the VDJdb data, the authors use 5-fold cross-validation instead of LOSO. Other evaluations were reported by using leave-one-out cross-validation of all unique TCRs (as defined by CDR3 sequence and V-gene). We compared ERGO when only the CDR3β sequence is utilized to the published TCRGP results for three specific human peptides from Dash et al.^17^ dataset. ERGO outperforms TCRGP models on most peptides, although ERGO was not trained to solve the SPB task for these specific peptides, but rather the more generic TPP task (Table 2, Table S5, figure 1D and Supp. Mat. methods for details of the training and test procedure for these and all other tests).

**Table 2:** Comparison between the different versions of the ERGO classifier (AE vs. LSTM and McPAS vs VDJdb) and existing classifier for the SPB task. Bolded values are the best results. The peptides here are the human peptides in Dash et al. dataset proposed by Jokinen et al. The other VDJdb peptides tested by the same authors are in the Supp. Mat. The values are the AUC over the test set of previousely unseen TCR for this peptide.

| Peptide | ERGO | | | | | | ERGO<br>best | TCRGP<br>($\beta$,3),<br>LOSO | TCRGP<br>($\beta$,3),<br>unique<br>LOO | TCRdist<br>($\beta$,3) |
| --- | --- | --- | --- | --- | --- | --- | --- | --- | --- | --- |
|  | McPAS |  | VDJdb |  | McPAS+<br>VDJdb |  |  |  |  |  |
|  | AE | LSTM | AE | LSTM | AE | LSTM |  |  |  |  |
| GLCTLVAML | 0.803 | 0.816 | 0.764 | 0.770 | 0.708 | 0.686 | 0.816 | 0.782 | **0.852** | 0.846 |
| NLVPMVATV | 0.835 | 0.821 | 0.665 | 0.686 | 0.624 | 0.632 | **0.835** | 0.587 | 0.651 | 0.680 |
| GILGFVFTL | 0.843 | 0.832 | 0.820 | 0.817 | 0.725 | 0.712 | **0.843** | 0.818 | 0.822 | 0.828 |

We used two-sample-logos^26^ to compare the CDR3 sequences of cognate TCRs for the three human peptides from Dash et al. dataset with TCRs that do not bind these peptides in the McPAS database (Figure 1E-G). Only 13 AA long TCRs were compared to avoid any alignment bias. While one can clearly see that different peptides have different signatures, it is interesting to see that the signature is not equally positioned among peptides. For the GLCTLVAML peptide, a signature is divided equally along the TCR, with a strong bias for initial CSA and not CAS. The NLVPMVATV signature is distributed following the standard CAS to the end of the CDR3, while the GILGFVFTL binding peptides are characterized by a dominant RS at position 6-7

The single peptide binding task can be extended to the single antigen protein task, where we predict whether a TCR would bind any peptide from a protein. Instead of testing whether an unseen TCR can bind a specific peptide, we tested whether it can bind any peptide from a target protein. The performance on this task varies drastically among target peptides, with AUC ranging from 0.71 to 0.97 (Table 3). This difference is not directly related to the number of target’s TCRs in the training set, but may rather represent the contribution of other factors not incorporated here, such as the alpha chain or the MHC.

**Table 3:**
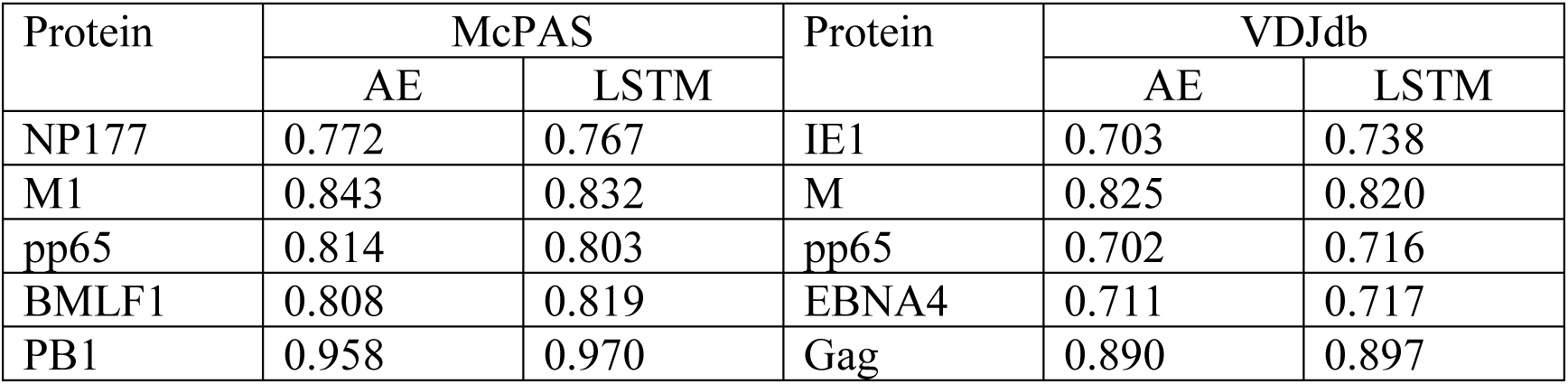
Comparison between the different versions of the ERGO classifier (AE vs. LSTM and McPAS vs VDJdb) for the binding to a specific antigen. There are no previous results on this task.

| Protein | McPAS |  | Protein | VDJdb |  |
| --- | --- | --- | --- | --- | --- |
|  | AE | LSTM |  | AE | LSTM |
| NP177 | 0.772 | 0.767 | IE1 | 0.703 | 0.738 |
| M1 | 0.843 | 0.832 | M | 0.825 | 0.820 |
| pp65 | 0.814 | 0.803 | pp65 | 0.702 | 0.716 |
| BMLF1 | 0.808 | 0.819 | EBNA4 | 0.711 | 0.717 |
| PB1 | 0.958 | 0.970 | Gag | 0.890 | 0.897 |

### Determining the target of a TCR (MPS task)

To use a TCR as a useful biomarker, one should be able to predict which specific peptide it binds. To test for that, we computed the accuracy (as measured by the sum of the diagonal in the confusion matrix) of predicting the proper target, with a different number of possible targets (Figure 2A). Again, ERGO was not trained for this task, but for the TPP task. The targets were the peptides with the highest number of binding TCR in the databases (Table S6). The AE produces better accuracies than the LSTM and the prediction for the AE and VDJdb yields better accuracies than McPAS. An important result is that the accuracy plateaus at 0.5 even for 10 peptides, suggesting that high accuracy can be obtained even when choosing from a large number of peptides.

**Figure 2:**
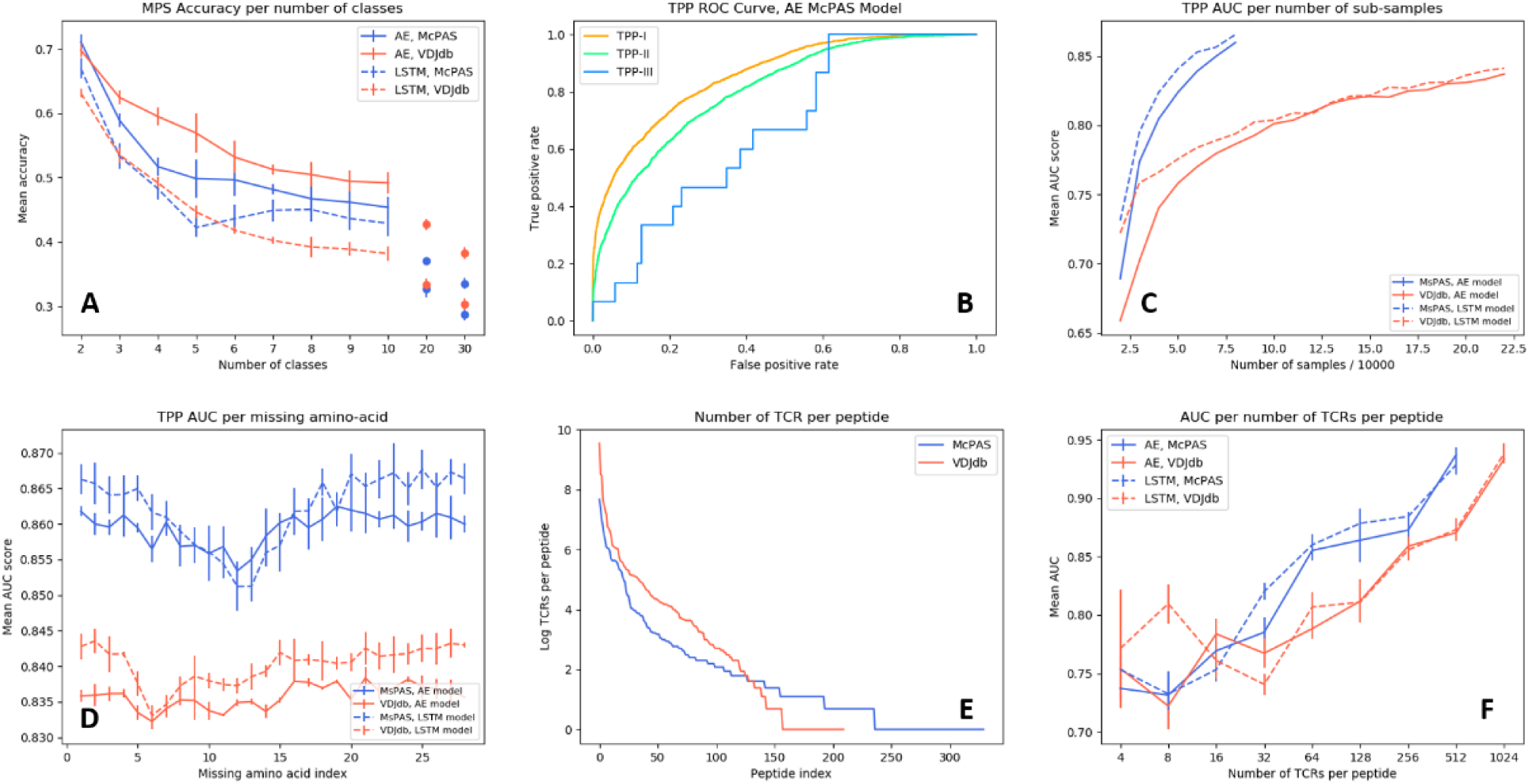
A) AUC per number of TCR-peptide pairs in McPAS-TCR and VDJdb datasets. B) ROC curve of TPP-I, II and III AE models performance on McPAS dataset C) AUC for TPP-I as a function of sub-sample size. D) AUC of TPP-I per missing amino-acids index. E) Number of TCRs per peptide distribution in McPAS-TCR and VDJdb datasets, logarithmic scale. F) AUC per number of TCRs per peptide bins (bins are union of all TCRs that match peptides with total number of TCRs in a specific range).

### Distinguishing TCRs binding different targets (TPP task)

A more important task from a diagnostic point of view would be to distinguish between TCRs binding different peptides for any set of either known or previousely unseen TCRs and peptides. To test the specificity of the prediction, we evaluated ERGO’s AUC on the three TPP tasks. The easiest task (TPP-I) is predicting unknown TCR-peptide parings (AE AUC value 0.86). A more complex task is the prediction of pairs containing a known peptide with an unknown TCR (TPP-II – AE AUC value 0.81). The hardest pairing task is to predict the binding of a previousely unseen peptide and a previousely TCR (TPP-III). This task has never been tested and reaches an AUC 0.669 (Table 4 and Figure 2B).

**Table 4:** AUC of TPP task with either known peptide and TCR (but unknown pairing TPP-I), known peptide unseen TCR (TPP-II) and unseen peptide and TCR (TPP-III). The results are the test AUC using either AE or LSTM on McPAS and VDJdb separately or on the joined dataset. The final column is the prediction of tumor antigens TCR binding. Again the AE consistently outperforms the LSTM, except for the TPP-III task, where increasing the training set size and the complexity of the encoders improves the performance.

| Evaluation AUC | McPAS |  | VDJdb |  | McPAS+ VDJdb |  | Tumor |  |
| --- | --- | --- | --- | --- | --- | --- | --- | --- |
|  | AE | LSTM | AE | LSTM | AE | LSTM | AE | LSTM |
| TPP-I | **0.860** | 0.859 | 0.840 | 0.842 | 0.776 | 0.761 | 0.805 | 0.813 |
| TPP-II | **0.810** | 0.798 | 0.792 | 0.764 | 0.770 | 0.745 | 0.805 | 0.813 |
| TPP-III | 0.601 | 0.562 | 0.669 | 0.522 | 0.636 | **0.674** | 0.570 | 0.646 |

To test if this performance can be improved by enlarging the training set to better learn the generic properties, we trained ERGO on McPAS and VDJdb simultaneously, and indeed the more complex LSTM encoder reached a higher AUC of 0.674, suggesting that further increasing the training set would improve the accuracy.

To further test the effect of the training set size, we subsampled the training set and tested the TPP-I AUC score for different sample sizes. The AUC increased with sample size and did not seem to saturate at the current sample size (Figure 2C). Some peptides have many reported TCRs binding them, while some have a single reported binding TCR. (Figure 2E) We tested whether a larger number of reported binding TCR improves accuracy (Figure 2F). Again, a higher number of bound TCRs induces higher prediction AUC, suggesting that larger datasets would further improve ERGO’s performance.

### Prediction of TCR-neoantigen binding

ERGO may be essential for the development of future TCR-based diagnostic tools. However, it can already be used for the detection of TCRs that bind specific tumor antigens. Given a neoantigen extracted from full genome sequencing of tumors^27,28^ and a target TCR, one could estimate the binding probability of the TCR to such a neoantigen. To test for that, we applied ERGO to neoantigen binding prediction; We used a positive dataset of cancer neoantigen peptides and their matching TCRs, published by Zhang et al.^29^, and expanded it with TCR-matching neoantigens in the McPAS-TCR and VDJdb databases. We tested again TPP-I, TPP-II and TPP-III. (Table 4), and got a high AUC for TPP-I and II (above 0.8), and 0.65 for the most complex TPP-III task. A caveat of this analysis may be that it was performed on a comparison of a dataset of TCRs binding neo-antigens and T cells from repertoires of healthy donors. Thus, formally this is not a direct measurement of the possibility of detecting neo-antigen specific TCRs within a donor.

### Comparison of TPP with literature

While TPP-III was never previously tested, TPP-II was recently tested by Jurtz et al.^19^, who used a convolutional neural network (CNN) based model, NetTCR, for predicting binding-probabilities of TCR-HLA-A*02:01 restricted peptide pairs. An IEDB dataset was used to train the model. The MIRA assay provided by Klinger et al.^30^ was used for evaluating the model by testing the model performance on shared IEDB and MIRA peptides and new TCRs. Jurtz et al. used two models in their experiments. One was trained with positive IEDB examples and only negative examples made from the IEDB dataset itself (no additional sources) while another model had also additional naïve negatives^31^. We used the united IEDB and MIRA dataset provided by Jurtz et al. and created also negative examples from that dataset. We trained ERGO models with 80% of the united data (positive and negative examples) and evaluated the model performance on the rest of the data (20%). Again, ERGO significantly outperformed the current results, 0.88 vs 0.73 (Table S3).

### CDR3 sequence characteristics

To test which position along the CDR3 has the strongest effect on the binding prediction, we trained ERGO ignoring one TCR amino-acid position at a time, by nullifying the position in the autoencoder based model or by skipping that position input in the LSTM based model (Figure 2D). Omitting each one of the central amino-acids of the TCR’s CDR3 beta (positions 7 to 15) impairs the model’s performance, especially in the LSTM-based model. The autoencoder-based model is more stable than the LSTM based model, perhaps due to exposure to a variety of TCRs in the TCR autoencoder pre-training.

## Discussion

We propose a set of standard tests to evaluate the accuracy of TCR-peptide binding and show that training a model using a combination of deep learning methods and curated datasets on the complex task of pairing random peptides and TCR can lead to high accuracy on all other tests. The main element affecting prediction accuracy is the training size. Enlarging the database improves the prediction accuracy for unseen peptides. In addition, when subsampling the existing datasets, the accuracy increases with sample size and does not seem to saturate at current sample size (Figure 2C).

Several other elements can affect the results, such as the V and J gene used and the alpha chain. In general, TCR-sequencing has often been limited to the TCR β chain due to its greater combinatorial and junctional diversity^10^ and to the fact that a single TCRβ chain can be paired with multiple TCRα chains^32^. Pogorelyy et al.^33^ have shown concordance between TCRα and TCRβ chain frequencies specific for a given epitope and suggested this justifies the exclusive use of TCRβ sequences in analyzing the antigen-specific landscape of heterodimeric TCRs. Currently, most proposed classifiers have used only CDR3 beta chains^4,34^ but some attempts have been made to include alpha chains^17^. Only recently, with single-cell techniques that enable pairing of α and β chains sequences, more data on alpha-beta TCRs is accumulating^35^. Once large-scale curated alpha-beta TCR-peptide datasets are available, their integration into the current method is straight forward.

ERGO is based on LSTM networks to encode sequential data. Previous models by Jurtz et al.^19^ used convolutional neural networks (CNN) for a similar task. While CNN are good at extracting position-invariant features, RNN (in particular LSTM) can catch a global representation of a sequence, in various NLP tasks^36^. Similarly, we did not use attention-based models^37^ since the TCR can bind the peptide MHC at different angles and specific TCR positions are not well correlated with specific peptide positions^38^.

ERGO randomly initializes our amino-acid embeddings and trains the embeddings with the model parameters. Using word-embedding algorithms such as Word2Vec^39^ or GloVe^40^ can give a good starting point to the embeddings. Special options for amino-acids pre-trained embeddings include the use of BLOSUM matrix^41^ or Kidera-factors-based manipulations^42^. As pre-trained embedding usually provides better model results, we plan to further test such encodings.

The prediction method presented here can serve as a first step in identifying neoantigen-reactive T cells for adoptive cell transfer (ACT) of tumor-infiltrating lymphocytes (TILs) targeting neoantigens^43^. The ERGO algorithm can accelerate the preliminary selection of valid target epitopes and corresponding TCRs for adoptive cell transfer. Finally, an important future implication would be to predict TCR-MHC binding, such prediction can be crucial for improving mismatched bone marrow transplants^44^.

## Acknowledgments

We wish to thank Prof. Luning Prak for helpful critiques and suggestions.

IS developed the formalism and implemented it.

HB designed the initial formalism.

YL supervised the work and wrote the manuscript.

NT developed the libraries and wrote the manuscript.

SD designed the TCR autoencoder formalism.

## Notes

#### Summary of Updates

New tests are proposed and the dataset is extended. Previous tests were removed for consistency with literature.

https://github.com/louzounlab/ERGO

