## Supplementary Materials for "Prediction of specific TCR-peptide binding from large dictionaries of TCR-peptide pairs"

### Materials and Methods

##### The Data

Three TCR-peptide datasets were used in the attachment prediction task. McPAS-TCR dataset was downloaded from http://friedmanlab.weizmann.ac.il/McPAS-TCR/ and VDJdb dataset was downloaded from https://vdjdb.cdr3.net/. We used a dataset of cancer neoantigen peptides and their matching TCRs, published by Zhang et al.[30]. A set of cancerous peptides was made for extracting TCRs matching to these peptides also in McPAS-TCR and VDJdb databases. We extended the original cancer dataset to include all TCRs-cancerous peptide pairs in all datasets. The data were processed into TCR-peptide pair files, using only TCRβ chains and valid TCR and peptide sequences.

The TCR autoencoder was trained on a data which was derived from a prospective clinical study (NCT00809276) by Kanakry et al.[44] The dataset is freely available at the Adaptive database (www.adaptivebiotech.com) that provides open access to a variety of datasets of TCRs next generation sequencing.

##### Datasets studied

In each model, training data was loaded as batches of positive and negative examples. For the positive examples, we took the existing TCR-peptide pairs in the database and split it to a train set and a test set. For creating the negative examples for the TPP-I task, we first chose a peptide randomly from the peptides in the training set. Then, we chose five random TCRs from the training set that are not reported to bind this peptide, to create five internal wrong pairs. A similar process was done to create a test set containing positive and negative examples. Thus, the number of negative examples is 5 times larger than the number of positive examples in both train and test sets.

#### The Models

We used two models for predicting TCR-peptide binding. The models use deep-learning architectures to encode the TCR and the peptide. Then the encodings are fed into multilinear perceptron (MLP) to predict the binding probability. Two encoding methods are applied – LSTM acceptor encoding and Autoencoder-based encoding. The peptide is always encoded using the LSTM acceptor method, so the two models differ in the TCR encoding method.

##### LSTM Acceptor

First, the amino acids were embedded using an embedding matrix. We set each amino acid an embedding vector, randomly initialized. Next, the TCR or the peptide was fed into a LSTM network as a sequence of vectors. The LSTM network outputs a vector for every prefix of the sequence; we used the last output as the encoding of the whole sequence. We used two different embedding matrices and LSTM parameters for the TCRs and the peptides encodings. The embedding dimension of the amino acids was 10. We use two-layered stacked LSTM, with 500 units at each layer. A dropout rate of 0.1 was set between the layers.

**TCR Autoencoder**

The TCR autoencoder was trained before training the Autoencoder-based attachment prediction model. In order to train the TCR autoencoder, first we added a 'stop-codon' at the end of every TCR CDR3 sequence. Each amino acid was represented as one-hot vector of 21 numbers (20 possible amino acids and an additional stop codon) where all values were set to zeros except one index of the corresponding amino acid which was set to 1. Each of the CDR3 vector representations one-hot vectors were joined and, terminated with a 'stop codon' one-hot vector. Zero padding was then added to the CDR3 vectors, completing the vectors to the maximum lengths chosen according to the data lengths distribution. Each zero codon was represented as fully zeroed one-hot vector.

The concatenated TCR vectors were fed into the Autoencoder network, which was based on a combination of linear layers, creating a similar 'encoder' and 'decoder' networks. In the encoder the TCRs were first put into a layer with 300 units, then into a layer with 100 units, and then into the encoding layer with 100 units. This layer output was used to encode the TCR in the trained autoencoder model. We used Exponential Linear Unit (ELU) activation between the linear layers and dropout with rate of 0.1. The decoding layers were similar to the encoding layers in the reverse order – first the encoded TCR vectors were fed into a layer with 100 units, then into a layer with 300 units, and then into a layer with the original TCR concatenated one-hot vector length units. We used softmax on the last decoder layer output on every sub-vector matching to an input amino acid one-hot vector position.

We used Mean Squared Error (MSE) loss (when the decoder output should be like the concatenated one-hot input). The autoencoder was trained using Adam optimizer with learning rate of 1e-4, We used batching with batch size 50. The autoencoder was trained for 300 epochs.

In order to read the TCR from the decoding vector, first we split the long vector into 'one-hot' like vectors. We back-translated the one-hot vectors into amino-acids by taking the amino acid matching to the maximal value index in the vector (which should be 1). We dropped all amino acids from the stop codon and forward to get a sequence of amino acid which should be the TCR.

The autoencoder was trained with 80% of the data and was evaluated with the rest of it.

The accuracy of the autoencoder was tested using the predictions of the test set. The number of mismatches between a CDR3 input vector to its prediction after trimming both vectors according to the start position of the codon vector in the predicted representation, removing the stop vector along with the zeros chain, leaving only the CDR3 representation to be accurate. Three types of accuracies were considered in this project, considering not only exact match as true prediction, but also allowing for one or two mismatches (Table S1).

***Table S1:*** *TCR autoencoder accuracy per number of mismatches allowed in sequence decoding.*

| Number of mismatches allowed | Autoencoder Accuracy |
|---|---|
| 0 | 0.920 |
| 1 | 0.982 |
| 2 | 0.993 |

**MLP Classifier (also mentioned as FFN)**

In both models, the TCR encoding was concatenated to the peptide encoding and fed into the MLP. The MLP contains one hidden layer with as units as half of the concatenated vector size and sigmoid is used on the output of the last layer to get a probability value. In both models the activation in the MLP is Leaky ReLU. Dropout with rate of 0.1 was set between layers.

**Model configurations**

As mentioned, we used two models, the LSTM based model and the autoencoder based model. We trained the embeddings, the LSTM parameters and the MLP in the first model, and the TCR autoencoder, peptide LSTM encoder and MLP parameters in the second model. The trained TCR autoencoder parameters were loaded to the autoencoder based model and are trained again within all model parameters.

We used Binary Cross Entropy (BCE) loss. Since we get 5 times more negative samples than positive samples according to the described sampling method, the loss is weighted respectively by a factor of 5/6 for positive samples and by 1/6 for negative samples. The optimizer was Adam with learning rate of 1e-3 and weight decay 1e-5. We used batching with batch size 50. The model was trained for 100 epochs. The models used 80% of the data for training and 20% for evaluation for all datasets.
All models were implemented with PyTorch library in Python programming language. The prediction models were evaluated using Area Under the Curve (AUC) score.

**Hyperparameters Tuning**

Both LSTM based model and the Autoencoder based model hyperparameters were optimized using a grid search in the hyperparameters space. The hyperparameters to optimize were the embedding matrix dimension, the LSTM dimensions, learning rate, weight decay, activation functions etc. All models were tested with the same grid search.

#### Experiments configuration

At the broad level, the ERGO model was trained and designed to solve the TPP-I task. Since the train and the test set are chosen randomly for each training process (as described above), 5 trained models along with their matching train and test set were analyzed, for each database (McPAS or VDJdb) and model type (LSTM based or AE based). Train and test sizes are detailed in Table S2.

***Table S2:*** *Train and test sizes of all trained models for the different versions of the ERGO classifier (AE vs. LSTM and McPAS vs VDJdb).*

| **Iteration** | **Database** | **McPAS** | | | | **VDJdb** | | | |
|---|---|---|---|---|---|---|---|---|---|
| | **Model type** | **AE** | | **LSTM** | | **AE** | | **LSTM** | |
| | **Dataset** | **Train** | **Test** | **Train** | **Test** | **Train** | **Test** | **Train** | **Test** |
| 1 | **TCR/Peptide pairs** | 66985 | 17316 | 67646 | 16659 | 209713 | 52758 | 209779 | 52693 |
| | **Positive pairs** | 11164 | 2886 | 11274 | 2776 | 34952 | 8793 | 34963 | 8782 |
| | **Negative pairs** | 55821 | 14430 | 56372 | 13883 | 174761 | 43965 | 174816 | 43911 |
| | **New test TCRs** | | 2119 | | 2047 | | 5852 | | 5873 |
| | **New test peptides** | | 14 | | 28 | | 10 | | 5 |
| 2 | **TCR/Peptide pairs** | 67351 | 16952 | 67094 | 17210 | 211209 | 51264 | 209396 | 53078 |
| | **Positive pairs** | 11225 | 2825 | 11182 | 2868 | 35201 | 8544 | 34899 | 8846 |
| | **Negative pairs** | 56126 | 14127 | 55912 | 14342 | 176008 | 42720 | 174497 | 44232 |
| | **New test TCRs** | | 2092 | | 2118 | | 5734 | | 5918 |
| | **New test peptides** | | 21 | | 15 | | 14 | | 12 |
| 3 | **TCR/Peptide pairs** | 67286 | 17018 | 67592 | 16711 | 209538 | 52933 | 209832 | 52638 |
| | **Positive pairs** | 11214 | 2836 | 11265 | 2785 | 34923 | 8822 | 34972 | 8773 |
| | **Negative pairs** | 56072 | 14182 | 56327 | 13926 | 174615 | 44111 | 174860 | 43865 |
| | **New test TCRs** | | 2105 | | 2011 | | 5858 | | 5820 |
| | **New test peptides** | | 24 | | 17 | | 15 | | 16 |
| 4 | **TCR/Peptide pairs** | 67644 | 16658 | 67489 | 16814 | 210232 | 52245 | 210440 | 52035 |
| | **Positive pairs** | 11274 | 2776 | 11248 | 2802 | 35038 | 8707 | 35073 | 8672 |
| | **Negative pairs** | 56370 | 13882 | 56241 | 14012 | 175194 | 43538 | 175367 | 43363 |
| | **New test TCRs** | | 2076 | | 2064 | | 5804 | | 5803 |
| | **New test peptides** | | 25 | | 24 | | 9 | | 18 |
| 5 | **TCR/Peptide pairs** | 67332 | 16968 | 67014 | 17290 | 209670 | 52802 | 209365 | 53108 |
| | **Positive pairs** | 11222 | 2828 | 11169 | 2881 | 34945 | 8800 | 34894 | 8851 |
| | **Negative pairs** | 56110 | 14140 | 55845 | 14409 | 174725 | 44002 | 174471 | 44257 |
| | **New test TCRs** | | 2088 | | 2122 | | 5895 | | 5914 |
| | **New test peptides** | | 19 | | 20 | | 9 | | 13 |

#### Single peptide binding

For computing single peptide binding score, samples from each test set were observed. For every peptide we looked for the pairs in the test det containing that peptide (positive and negative samples). The ROC and AUC scores were computed according to the model prediction of those pairs. SPB scores were computed for the five frequent peptides in each database (Table 1) and three human peptides appearing

in Dash et al dataset (Table 2). Results for peptides with more than 50 reported binding TCRs are in the Supp Mat. Table S1. Mean AUC scores are reported.
Single protein scores are extracted in a similar way, by analyzing all pairs in the test set that contain a peptide of the specific protein.

**Multi-Peptide Selection**

At first, number of classes k was set. Trained model prediction scores were extracted for each TCR in the test set, paired with every peptide from the top k frequent peptides in the relevant database. The TCR target was predicted to be the peptide which got the maximal score as the pair complement. Accuracy was computed using the true samples in the test set. This was done for number of classes ranged between 2 and 10, as well as 20 and 30. Mean accuracies are shown in Figure 2A.

**TCR-Peptide Pairing**

New test TCRs and new test peptides were deducted from the train and test sets. TPP-I score is the AUC of the model predictions of the original test set. TPP-II is the AUC of the predictions of the new test TCR positive and negative samples. TPP-III is the AUC of the predictions of new test TCR and new test peptide pairs, positive and negative samples.

**Train data sub-sampling**

All models were trained and evaluated using the same train and test partition. Every model train set was a sub-sample of the original train set, while the test set remained the same. 10000 new train samples were added at each iteration.

**Missing positions training**

Again, all models were evaluated with the same test set and a train/test partition. In this experiment, the train data was modified by dropping a single amino acid in a specific position at a time. Practically, this was done be deleting this position for all TCRs in the LSTM based model, or by nullifying the relevant position in the one-hot encoding of the TCRs in the AE based model. This experiment was repeated five times, Mean TPP-I scores are shown in Figure 2D.

**TPP per number of TCRs per peptide**

First, TCR records per peptide were counted in the original McPAS and VDJdb databases. Given a test set, the test pairs were divided to bins, according to the number of TCR records per peptide in the original database. The differences between the bins were on exponential scale. AUC score was computed for each bin. Mean AUC scores are shown in Figure 2F.

**Comparison with NetTCR**

The united IEDB and MIRA dataset was downloaded from https://github.com/mnielLab/netTCR. Unfortunately, the authors did not publish the IEDB train data separated from the MIRA test data, thus we had to evaluate ERGO in another train/test partition. We used 80% of the IEDB and MIRA data for training and the rest of it (20%) for testing. Additional 'C' prefix and 'F' suffix were added to each TCR sequence. The MIRA data was containing new test TCRs (but was not evaluated with new test peptides), therefore we compare NetTCR results with ERGO TPP-II scores (Table S3).

***Table S3:*** *Comparison between the different versions of the ERGO classifier (AE vs. LSTM) and existing NetTCR models. TPP-II results are shown.*

| **Model** | **Train data** | **Test data** | **TPP-II score** |
|---|---|---|---|
| ERGO AE | IEDB + MIRA (80%) | IEDB + MIRA (20%) | 0.886 |
| ERGO LSTM | IEDB + MIRA (80%) | IEDB + MIRA (20%) | 0.883 |
| NetTCR INT_NEG | IEDB | MIRA (shared IEDB peptides) | 0.697 |
| NetTCR ADD_NEG | IEDB + additional negatives | MIRA (shared IEDB peptides) | 0.727 |

***Table S4:*** *Comparison between the different versions of the ERGO classifier (AE vs. LSTM and McPAS vs VDJdb) for the SPB task. Peptides with more than 50 TCRs in each database are shown. The values are the AUC over the test set of unseen TCR for this peptide.*

| **McPAS** | | | **VDJdb** | | |
|---|---|---|---|---|---|
| **Model** / **Peptide** | **AE** | **LSTM** | **Model** / **Peptide** | **AE** | **LSTM** |
| LPRRSGAAGA | 0.772431 | 0.767037 | KLGGALQAK | 0.695061 | 0.731208 |
| GILGFVFTL | 0.843091 | 0.832766 | GILGFVFTL | 0.820975 | 0.817835 |
| NLVPMVATV | 0.835317 | 0.821623 | NLVPMVATV | 0.665098 | 0.686264 |
| GLCTLVAML | 0.803314 | 0.816092 | AVFDRKSDAK | 0.676548 | 0.695028 |
| SSYRRPVGI | 0.969659 | 0.980026 | RAKFKQLL | 0.828297 | 0.825514 |
| RFYKTLRAEQASQ | 0.967792 | 0.936695 | ELAGIGILTV | 0.735488 | 0.862051 |
| SSLENFRAYV | 0.942426 | 0.942475 | GLCTLVAML | 0.764231 | 0.770183 |
| WEDLFCDESLSSPEPPSSSE | 0.901104 | 0.920351 | IVTDFSVIK | 0.763487 | 0.764299 |
| CRVLCCYVL | 0.770348 | 0.768169 | SSYRRPVGI | 0.989022 | 0.986433 |
| ASNENMETM | 0.940957 | 0.944221 | SSLENFRAYV | 0.969169 | 0.970508 |
| ELAGIGILTV | 0.828654 | 0.816382 | RLRAEAQVK | 0.726325 | 0.695372 |
| LLWNGPMAV | 0.856099 | 0.838728 | TTPESANL | 0.966307 | 0.970118 |
| VTEHDTLLY | 0.810553 | 0.815736 | CTPYDINQM | 0.959594 | 0.968356 |
| TPRVTGGGAM | 0.803075 | 0.787879 | LLWNGPMAV | 0.725481 | 0.75429 |
| HGIRNASFI | 0.959277 | 0.959055 | HGIRNASFI | 0.971845 | 0.981121 |
| EAAGIGILTV | 0.854633 | 0.85533 | ASNENMETM | 0.974924 | 0.976743 |
| FRCPRRFCF | 0.824175 | 0.790418 | PKYVKQNTLKLAT | 0.691382 | 0.663138 |
| VEALYLVCG | 0.907081 | 0.935765 | FRDYVDRFYKTLRAEQASQE | 0.952822 | 0.936532 |
| LSLRNPILV | 0.902143 | 0.924408 | KRWIILGLNK | 0.842182 | 0.837442 |
| RPHERNGFTVL | 0.714755 | 0.731987 | SSPPMFRV | 0.984529 | 0.985437 |
| SSPPMFRV | 0.976377 | 0.977021 | STPESANL | 0.969857 | 0.974328 |
| RAKFKQLL | 0.832942 | 0.814849 | LSLRNPILV | 0.93783 | 0.963003 |
| KAFSPEVIPMF | 0.843734 | 0.913372 | CINGVCWTV | 0.813524 | 0.827376 |
| NLNCCSVPV | 0.850514 | 0.789345 | TPRVTGGGAM | 0.845308 | 0.841128 |
| MEVGWYRSPFSRVVHLYRNGK | 0.987286 | 0.983272 | LLLGIGILV | 0.728453 | 0.682906 |
| KRWIILGLNK | 0.767641 | 0.766451 | KAFSPEVIPMF | 0.840425 | 0.902281 |
| FPRPWLHGL | 0.795516 | 0.884843 | ATDALMTGY | 0.884728 | 0.83388 |
| TVYGFCLL | 0.874119 | 0.911252 | VTEHDTLLY | 0.625833 | 0.639702 |
| KMVAVFYTT | 0.757869 | 0.79095 | EIYKRWII | 0.775054 | 0.768612 |
| RPRGEVRFL | 0.862574 | 0.922006 | FLKEKGGL | 0.738713 | 0.747265 |
| ATDALMTGY | 0.941682 | 0.869135 | GTSGSPIVNR | 0.84617 | 0.854691 |
| YVLDHLIVV | 0.878755 | 0.7948 | TVYGFCLL | 0.970757 | 0.972494 |
| VVLSWAPPV | 0.710898 | 0.721487 | GTSGSPIINR | 0.862618 | 0.864304 |
| YSEHPTFTSQY | 0.868444 | 0.719544 | SQLLNAKYL | 0.987209 | 0.985011 |
| HPKVSSEVHI | 0.840526 | 0.846167 | LPRRSGAAGA | 0.619644 | 0.585148 |
| | | | NLSALGIFST | 0.707337 | 0.756388 |

| | | | | | |
|---|---|---|---|---|---|
| | | | RPRGEVRFL | 0.89807 | 0.951702 |
| | | | ARMILMTHF | 0.920818 | 0.9468 |
| | | | SFHSLHLLF | 0.736924 | 0.7445 |
| | | | QARQMVQAMRTIGTHP | 0.650485 | 0.665243 |
| | | | KAVYNFATC | 0.894006 | 0.932316 |
| | | | GLIYNRMGAVTTEV | 0.642705 | 0.606565 |
| | | | TPQDLNTML | 0.752846 | 0.737685 |
| | | | IPSINVHHY | 0.75858 | 0.74078 |
| | | | DPFRLLQNSQVFS | 0.67448 | 0.747417 |
| | | | YVLDHLIVV | 0.678381 | 0.723087 |
| | | | YSEHPTFTSQY | 0.802649 | 0.869689 |
| | | | KLVALGINAV | 0.634441 | 0.718725 |
| | | | RTLNAWVKV | 0.555511 | 0.672836 |
| | | | FLRGRAYGL | 0.90147 | 0.843084 |
| | | | FPRPWLHGL | 0.865248 | 0.760968 |
| | | | AMFWSVPTV | 0.591343 | 0.707997 |
| | | | TPGPGVRYPL | 0.797351 | 0.747012 |
| | | | KRWIIMGLNK | 0.853095 | 0.828291 |
| | | | AYAQKIFKI | 0.598942 | 0.595577 |
| | | | SLFNTVATLY | 0.677828 | 0.682458 |
| | | | FLYALALLL | 0.878 | 0.883727 |
| | | | GPGHKARVL | 0.544828 | 0.742347 |
| | | | LLFGYPVYV | 0.614185 | 0.666065 |
| | | | MLNIPSINV | 0.583162 | 0.635572 |
| | | | SLYNTVATL | 0.795793 | 0.77279 |
| | | | NEGVKAAW | 0.817963 | 0.724294 |
| | | | FLASKIGRLV | 0.669265 | 0.721543 |
| | | | QVPLRPMTYK | 0.821861 | 0.849146 |
| | | | SGPLKAEIAQRLED | 0.746869 | 0.631266 |
| | | | EPLPQGQLTAY | 0.880626 | 0.85187 |
| | | | SYIGSINNI | 0.965756 | 0.954853 |
| | | | HPVGEADYFEY | 0.94127 | 0.89122 |
| | | | FLYNLLTRV | 0.620185 | 0.708811 |
| | | | NAITNAKII | 0.968216 | 0.938254 |
| | | | GTSGSPIIDK | 0.72907 | 0.733715 |
| | | | ISPRTLNAW | 0.686906 | 0.722699 |
| | | | HPKVSSEVHI | 0.847606 | 0.73928 |
| | | | RPPIFIRRL | 0.803643 | 0.822873 |
| | | | LLDFVRFMGV | 0.773148 | 0.634493 |
| | | | QYDPVAALF | 0.617483 | 0.624344 |

***Supp. Mat. Table S5:*** *Comparison between the different versions of the ERGO classifier (AE vs. LSTM and McPAS vs VDJdb) and existing classifier for the SPB task. The peptides are the VDJdb peptides tested by Jokinen et al. The values are the AUC over the test set of unseen TCR for this peptide.*

| **Peptide** | **ERGO** | | **TCRGP (β,3)** | |
|---|---|---|---|---|
| | **AE** | **LSTM** | **LOSO** | **unique LOO** |
| IPSINVHHY | 0.758 | 0.74 | 0.852 | 0.797 |
| TPRVTGGGAM | 0.845 | 0.841 | 0.892 | 0.768 |
| NLVPMVATV | 0.665 | 0.686 | 0.912 | 0.838 |
| GLCTLVAML | 0.764 | 0.77 | 0.926 | 0.782 |
| RAKFKQLL | 0.828 | 0.825 | 0.887 | 0.729 |
| YVLDHLIVV | 0.678 | 0.723 | 0.682 | 0.541 |
| GILGFVFTL | 0.82 | 0.817 | 0.881 | 0.804 |
| PKYVKQNTLKLAT | 0.691 | 0.663 | 0.706 | 0.563 |
| CINGVCWTV | 0.813 | 0.827 | 0.819 | 0.887 |
| KLVALGINAV | 0.634 | 0.718 | 0.695 | 0.484 |
| ATDALMTGY | 0.884 | 0.833 | 0.678 | 0.677 |
| RPRGEVRFL | 0.897 | 0.951 | 0.801 | 0.694 |
| LLWNGPMAV | 0.725 | 0.754 | 0.825 | 0.813 |
| GTSGSPIVNR | 0.846 | 0.854 | 0.864 | 0.832 |
| GTSGSPIINR | 0.855 | 0.864 | 0.734 | 0.736 |
| KAFSPEVIPMF | 0.84 | 0.902 | 0.769 | 0.755 |
| TPQDLNTML | 0.752 | 0.737 | 0.798 | 0.742 |
| EIYKRWII | 0.775 | 0.768 | 0.75 | 0.904 |
| KRWIILGLNK | 0.839 | 0.837 | 0.702 | 0.535 |
| FRDYVDRFYKTLRAEQASQE | 0.952 | 0.936 | 0.893 | 0.822 |
| GPGHKARVL | 0.544 | 0.742 | 0.838 | 0.803 |
| FLKEKGGL | 0.738 | 0.747 | 0.817 | 0.766 |

***Supp. Mat. Table S6****: The peptides with the highest number of binding TCR in McPAS and VDJdb databases, in decreasing order. 30 most frequent peptides in both databases are shown.*

| **McPAS** | | **VDJdb** | |
|---|---|---|---|
| **Peptide** | **TCRs** | **Peptide** | **TCRs** |
| LPRRSGAAGA | 2145 | KLGGALQAK | 27842 |
| GILGFVFTL | 1920 | GILGFVFTL | 7008 |
| NLVPMVATV | 1134 | NLVPMVATV | 4991 |
| GLCTLVAML | 1091 | AVFDRKSDAK | 3534 |
| SSYRRPVGI | 653 | RAKFKQLL | 2720 |
| RFYKTLRAEQASQ | 602 | ELAGIGILTV | 2519 |
| SSLENFRAYV | 549 | GLCTLVAML | 1599 |
| WEDLFCDESLSSPEPPSSSE | 477 | IVTDFSVIK | 1480 |
| CRVLCCYVL | 435 | SSYRRPVGI | 1306 |
| ASNENMETM | 427 | SSLENFRAYV | 866 |
| ELAGIGILTV | 325 | RLRAEAQVK | 860 |
| LLWNGPMAV | 307 | TTPESANL | 859 |
| VTEHDTLLY | 280 | CTPYDINQM | 814 |
| TPRVTGGGAM | 279 | LLWNGPMAV | 697 |
| HGIRNASFI | 279 | HGIRNASFI | 558 |
| EAAGIGILTV | 278 | ASNENMETM | 536 |
| FRCPRRFCF | 266 | PKYVKQNTLKLAT | 484 |
| VEALYLVCG | 207 | FRDYVDRFYKTLRAEQASQE | 471 |
| LSLRNPILV | 203 | KRWIILGLNK | 413 |
| RPHERNGFTVL | 191 | SSPPMFRV | 326 |
| SSPPMFRV | 163 | STPESANL | 309 |
| RAKFKQLL | 144 | LSLRNPILV | 290 |
| KAFSPEVIPMF | 143 | CINGVCWTV | 284 |
| NLNCCSVPV | 128 | TPRVTGGGAM | 273 |
| MEVGWYRSPFSRVVHLYRNGK | 128 | LLLGIGILV | 233 |
| KRWIILGLNK | 99 | KAFSPEVIPMF | 227 |
| FPRPWLHGL | 88 | ATDALMTGY | 207 |
| TVYGFCLL | 87 | VTEHDTLLY | 203 |
| KMVAVFYTT | 74 | EIYKRWII | 176 |
| RPRGEVRFL | 65 | FLKEKGGL | 175 |
